# Sex, fitness decline and recombination – Muller’s ratchet vs. Ohta’s ratchet

**DOI:** 10.1101/2020.08.06.240713

**Authors:** Yongsen Ruan, Haiyu Wang, Lingjie Zhang, Haijun Wen, Chung-I Wu

**Author notes:** Correspondence to: Haijun Wen, Chung-I Wu.

## Abstract

It is generally accepted that the absence of recombination reduces the efficacy of natural selection for, or against, mutations. A special case is Muller’s Ratchet (MR) whereby non-recombining genomes experience irreversible fitness decline due to the accumulation of deleterious mutations. MR has been a main hypothesis for sexual reproduction as well as many other biological phenomena. We now ask whether the fitness decline can indeed be stopped if an asexual population turns sexual to become recombining. The possible fitness decline under recombination is referred to as Ohta’s Ratchet (OR). In comparison, MR is more effective in driving fitness reduction than OR, but only in a restricted parameter space of mutation rate, population size and selection. Outside of this space, the two ratchets are equally effective or, alternatively, neither is sufficiently powerful. Furthermore, beneficial mutations can affect the population fitness, which may diverge between the two ratchets, but only in a small parameter space. Since recombination plays a limited role in driving fitness decline, the operation of MR could be far less common in nature than believed. A companion report (see Supplement) surveying the biological phenomena attributed to MR indeed suggests the alternative explanations to be generally more compelling.

## Introduction

Natural selection would operate most effectively when free recombination permits every gene be tested independently. Imagine a genome harboring many beneficial and deleterious mutations of equal strength. In the absence of recombination, selection could not operate on either type of mutations, which collectively would appear neutral. The interference among various actions of selection has been formalized as the Hill-Robertson effect (HR; (Hill and Robertson 1966; Felsenstein 1974; Charlesworth, et al. 2009)).

A special case of HR is Muller’s Ratchet (MR; (Muller 1964; Haigh 1978; McVean and Charlesworth 2000)). A ratchet is a mechanism of uni-directional movement. In MR, the fitness of individuals would continue to decline due to the accumulation of deleterious mutations. Most important, the absence of recombination reduces the efficacy of negative selection in removing deleterious mutations; for example, if individuals carrying different deleterious mutations are equally unfit, then none could be removed by natural selection. MR has been invoked to explain the evolution of many phenomena, including sexual reproduction (Muller 1964; Smith 1978; Kondrashov 1988). As the fitness of the population continues to decline, it is conceivable that sexual reproduction and recombination would reverse the fitness decline by generating mutation-free genomes. Under some conditions (Kondrashov 1984; Gillespie 1998), the fitness gain via recombination would be sufficient to compensate for the 2-fold cost associated with sexual reproduction. Other phenomena attributed to MR include the degeneracy of the Y chromosome and the evolution of selfing (Charlesworth and Charlesworth 2000; Bachtrog 2013).

In published studies, the emphasis is on the efficacy of MR in driving the fitness decline (Felsenstein 1974; Haigh 1978; Gordo and Charlesworth 2000b; Gordo and Charlesworth 2000a; Neher and Shraiman 2012; Neher 2013; McDonald, et al. 2016; Desai 2020; Leu, et al. 2020). In this study, we approach the issue somewhat differently. We compare the presence and absence of recombination in driving the fitness decline under the same input of deleterious mutations. The process of fitness decline under free recombination is referred to as Ohta’s ratchet (OR) in honor of Tomoko Ohta’s contributions to the studies of slightly deleterious mutations (Ohta 1973; Ohta 1976, 1987; Ohta 1992). There is little doubt that the fitness decline under MR should be faster than under OR. The issue is a quantitative one – Can recombination stop, or at least substantially slow down, the fitness decline such that there is a significant advantage in gaining the ability to recombine?

The operation of the ratchets, either MR or OR, will likely be reversed by beneficial mutations of sufficient strength. It is hence expected that the selection efficacy would be higher in the presence of recombination and OR can be more easily reversed. The issue is again a quantitative one. This study will investigate the theoretical aspects of the ratchets including the interferences between beneficial and deleterious mutations. A companion report (Wang et al.; Appended to the Supplement) will examine the empirical evidence for, and against, MR as the driving force of the evolution of the said phenomena.

### Mutation accumulation with free recombination

We now present a standard population genetic model with free recombination. For simplicity, we use a haploid model with *N* individuals. The mutation rate is *u* mutations per generation per haploid genome. The proportion of beneficial and deleterious mutations is *p* and *q*, respectively. The rest, 1 – *p* – *q*, is neutral. The average fitness effect of beneficial and deleterious mutations is, respectively, 1+*s*_*p*_ and 1−*s*_*q*_. There is no epistasis in fitness and all loci are freely recombining.

#### Fixation rate of mutations

The fixation probability for a mutant with a selective coefficient of *s* (where *s* = *s*_*p*_ or -*s*_*q*_) is

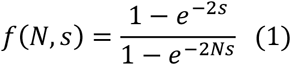

Then the relative fixation rates to neutral mutations for neutral, positive and negative mutations can be calculated as follows:

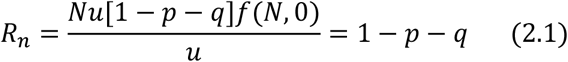

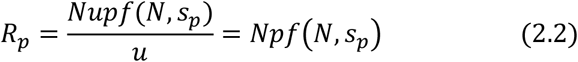

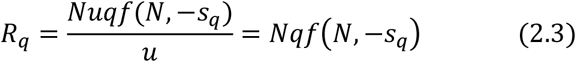

And the total fixation rate (equivalent to *Ka*/*Ks* or *dN*/*dS* (Spielman and Wilke 2015; Wu, et al. 2016)) is

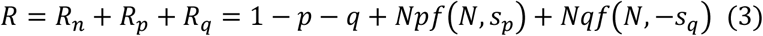

Now, the accumulation rate of fixed mutations is

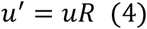

#### Fitness change as mutations accrue

Assuming multiplicative fitness values among all beneficial and deleterious mutations, we wish to know the conditions for the fitness to increase or decrease. Let *n*_*k*_ be the total number of fixed mutations. Then, the expected numbers of beneficial, deleterious and neutral mutations are

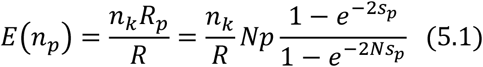

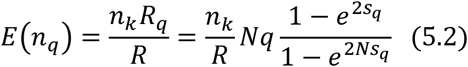

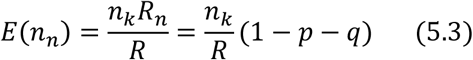

Let the *i*-th beneficial and *j*-th deleterious mutation have the selective coefficient of *s*_*pi*_ = *s*_*p*_, *s*_*qj*_ = *s*_*q*_. Then the expected fitness (multiplicative fitness) of an individual with *n*_*k*_ fixed mutation is

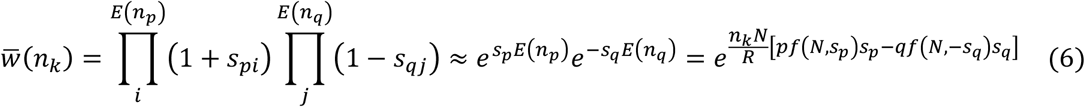

Starting from a population under the mutation-selection balance, the expect fixed mutation at generation *t* is *n*_*k*_(*t*) = *uRt*. And then the average fitness at generation *t* is

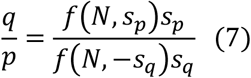

The fitness will decline when *pf*(*N, s*_*p*_)*s*_*p*_ < *qf*(*N*, −*s*_*q*_)*s*_*q*_ (i.e. 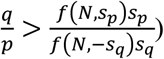), the average fitness of a sexual population will continue to decline over time. The opposite is true if the sign changes. When *pf*(*N*, *s*_*p*_)*s*_*p*_ = *qf*(*N*, −*s*_*q*_)*s*_*q*_, i.e.

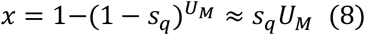

the average fitness stays constant over time.

### Fitness decline with deleterious mutations only

We will first consider a system with deleterious mutations only (*p* = 0 in equations above). In such a system, the population fitness will only go down in a ratchet-like mechanism. We will compare the rate of fitness decline with vs. without recombination. The influence of beneficial mutations will be analyzed in the next section.

#### The two ratchets – Muller’s vs. Ohta’s ratchet

The average fitness of the population would go down at different rates with or without recombination as illustrated in Fig. 1. As stated, we shall refer to the mechanism with recombination as Ohta’s ratchet (OR). The theory of OR, shown above, is standard textbook materials.

**Fig. 1.**
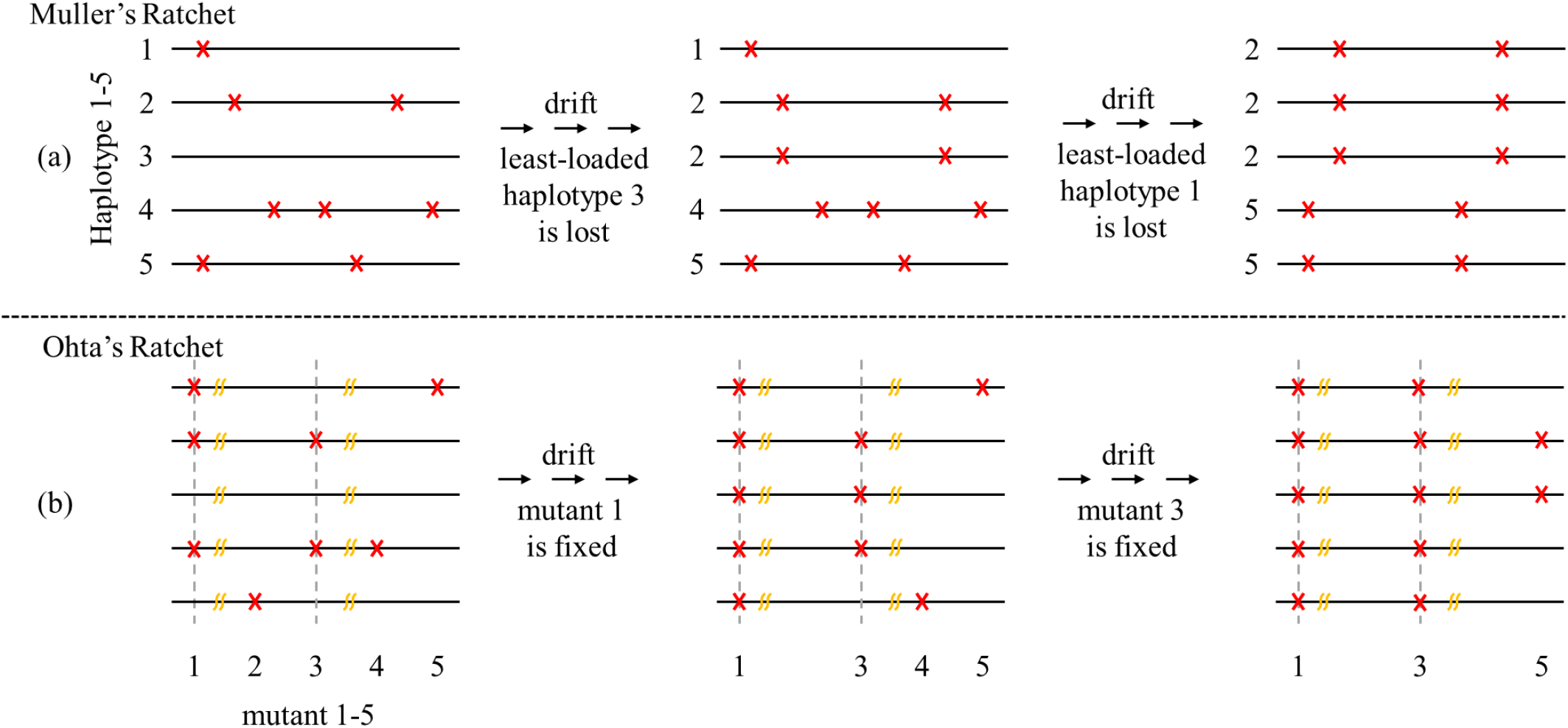
Accumulation of deleterious mutations by Muller’s ratchet (MR) and Ohta’s ratchet (OR). (a) In Muller’s ratchet, there is no recombination and the most fit haplotype would occasionally be lost due to genetic drift. Here, the fitness decline does not require fixation of mutations. (b) In Ohta’s ratchet, free recombination enables selection on each locus independently. The fitness decline becomes irreversible when the deleterious variant is fixed. Despite the differences, both processes operate most effectively in small populations with weak selection and high mutation rate.

In comparison, the mechanism of fitness decline without recombination is the classical Muller’s ratchet (MR; (Muller 1964; Felsenstein 1974; Haigh 1978; Gabriel, et al. 1993). Although the approximate formulae for the fitness decline of MR (Haigh 1978; Gessler 1995; Gordo and Charlesworth 2000a; Gordo and Charlesworth 2000b; Etheridge, et al. 2009; Neher and Shraiman 2012) and for the more general cases with beneficial mutations (Goyal, et al. 2012; Good, et al. 2014; Weissman and Hallatschek 2014) have been developed, the approximations are not always sufficiently accurate. In this study, we simulate a discrete-time Wright-Fisher model for the MR effect (see Methods).

#### Comparison of fitness under the two ratchets

Given a parameter set of (*N*, *s*_*q*_, *u*), the rate of mutation accumulation per generation will be denoted as *U*_*M*_ and *U*_*O*_, respectively for MR and OR. *U*_*M*_ is obtained by simulation as described in the Methods section and *U*_*O*_ = *uR* as shown in Eq. (4). Let *x* be the fitness reduction per generation under MR. After one generation, the fitness *w* decreases by *x*. Thus, 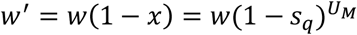 and

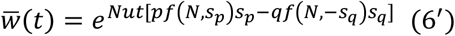

Similarly, the fitness reduction per generation for OR is

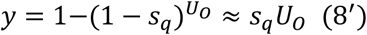

In the comparison between MR and OR, we use a series of values for *s*_*q*_(0.0001 – 0.1) and *N* (10 – 10,000). We let *u* = 0.03 which roughly corresponds to the per-generation rate for *Drosophila* (Haag-Liautard, et al. 2007; Keightley, et al. 2014) or *u* = 0.3 coding mutations per generation per gamete for human (Scally and Durbin 2012; Jonsson, et al. 2017; Ruan, et al. 2020). In Fig. 2a-2b where *Ns*_*q*_= 0.1 and *u* = 0.3, it is clear that MR and OR have similar rates of mutation accumulation and fitness decline. In Fig. 2c-2d where *Ns*_*q*_= 1 but *u* = 0.03, OR works more slowly. The time it takes to reduce the fitness by 50% increases by 2-fold (40,000 to 80,000 generations) from MR to OR. The gain in stalling the fitness decline by acquiring recombination seems rather small in these examples. It is hard to imagine the advantage of sexual reproduction when the two-fold cost associated with sex is taken into account (Smith 1978; Kondrashov 1984; Gillespie 1998).

**Fig. 2.**
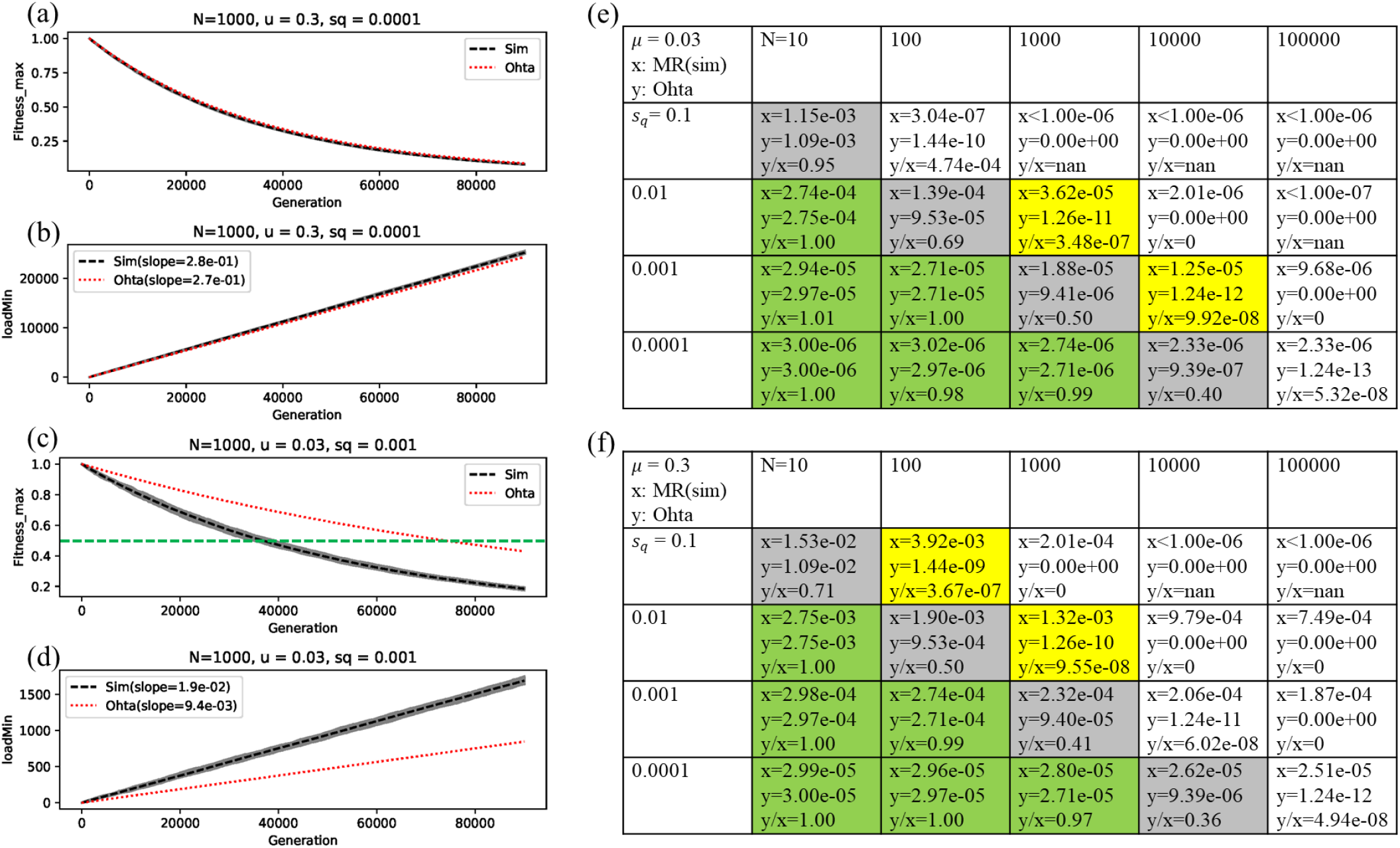
The relative efficacy of MR vs. OR with deleterious mutations only (*p*=0, *q*=1). (a, c) The fitness decline of the asexual population over time. (b, d) The number of mutations accrue over time in the same process. The black and red lines are, respectively, for MR and OR. The dynamics of OR is given in Eq. (6’). MR is done by computer simulations with 25 repeats (see Methods). In these two cases, the fitness decline happens with or without recombination. (e-f) Summary of fitness decline over a large range of parameter values in *u*, *sq* and *N*. The rate of fitness decline per generation is denoted *x* for MR and *y* for OR. The cases of *Ns*_*q*_<1 are shaded green whereby the two ratchets are equally effective. The cases of *Ns*_*q*_=1 are shaded grey whereby OR is also effective, albeit somewhat less than MR. In the cases without shading, OR generally does not operate but neither does MR operate fast enough to be biologically meaningful. Only the cases shaded yellow show MR, but OR, to work in a realistic time frame. This is the parameter space where recombination is biologically significant in preventing fitness decline (see text).

Fig. 2e and 2f present the results from the wider parameter space. We find that when *Ns*_*q*_ ≤ 0.1., the fitness dynamics of Muller’s ratchet is almost the same as that of Ohta’s ratchet (shaded green in Fig. 2). When *Ns*_*q*_ = 1, the speed of Ohta’s ratchet is about half of Muller’s ratchet (shaded grey). When *Ns*_*q*_ > 1, MR indeed moves faster than OR. However, in much of the parameter space (no shading), the faster movement of MR is in fact too slow to drive the evolution. The parameter sets whereby MR moves at a speed that seems biologically meaningful is shaded in yellow. For the lower mutation rate of *u* = 0.03, the speed is set at > 10^−5^ per generation and, for *u* = 0.3, the threshold is set at > 10^−3^. (We assume that small *u* is associated with short generation time and large population size; hence, a fitness loss of 10^−5^ is biologically significant. The opposite is assumed for species with larger *u*’s.) The speed is important because a slower ratchet can be easily reversed by factors that include beneficial mutations (see below and Wang et al. 2020).

In the analysis of MR vis-à-vis OR, the speed of MR at an intermediate rate may be biologically most significant (yellow highlight). Previous studies (Haigh 1978; Gordo and Charlesworth 2000b; Gordo and Charlesworth 2000a; Goyal, et al. 2012) have suggested that MR is most efficient when the number of individuals in the least-loaded (*n*_0_ = *Ne*^−*μ*/*s*_q_^) is less than 1. Since *n*_0_ for N = 100, *s*_*q*_=0.1, μ = 0.3 is 5, the contrast bolsters the argument that MR needs to be viewed by its efficiency relative to that of OR.

In short, the part of parameter space where MR works and works much better than OR is a tightly confined one. In other words, if the population experiences fitness decline, gaining the ability to recombine may not slow down the decline all that much. It is hard to see how the evolution of sex and a host of other major biological phenomena would have been driven by MR.

### Fitness change with both beneficial and deleterious mutations

The ratchet certainly could not operate indefinitely as beneficial mutations of a sufficient strength would reverse the fitness decline. For that reason, the ratchet, as well as the phenomena attributed to MR, must happen between the occurrences of such beneficial mutations. Here, we assume that all selection effects are multiplicative, as is usually done (Felsenstein 1974; Haigh 1978; Goyal, et al. 2012; Neher and Shraiman 2012; Shi, et al. 2019; Xu, et al. 2019). In reality, a beneficial mutation may annul the effects of many strongly deleterious mutations; for example, when the deleterious mutations are on the same pathway that is bypassed by the beneficial mutation (see Discussion). Nevertheless, for the sake of comparing the efficacy of MR and OR, multiplicative fitness may provide a glimpse of the power of recombination.

Eq. (7) defines the condition for beneficial mutations to reverse OR. If we assume *s*_*p*_ = *s*_*q*_ = 0.001 and *N* = 10,000 such that *Ns* = 10, then *p*/*q* of Eq. (7) would be ~ 2 × 10^−9^. In other words, beneficial mutations only need to be two billions as frequent as deleterious mutations to stop OR. It is expected that MR could be stopped by fewer and/or weaker beneficial mutations because the efficacy of selection is lower in the absence of recombination (Hill and Robertson 1966; Felsenstein 1974; Comeron, et al. 2008; Charlesworth, et al. 2009). Nevertheless, the asymmetry between beneficial and deleterious mutations in driving the change in population fitness is so great that recombination is not expected to override the asymmetry between beneficial and deleterious mutations.

Fig. 3 shows the decline in population fitness driven by MR vs. OR when *Ns*_*q*_= 1. The 8 panels differ in the relative strength of selection (*s*_*p*_/*s*_*q*_, row 1 to row 4) and the relative mutation frequency (*p*/*q*, column 1 and column 2). In panels a to c, the fitness decline would happen by both MR and OR although the speed is higher by MR. In panels f to h, the fitness would increase by either mechanism, but faster by OR. There is a small parameter space where the fitness would decline by MR and would increase by OR (see Fig. 3d and 3e). As the frequency of beneficial mutations become higher (Fig. 3f vs. 3e), both MR and OR would be reversed. Most important, when *Ns*_*p*_ increase to 10, neither process would operate (unless *p*/*q* is fairly close to 0). Note that *Ns*_*p*_ = 10 indicates rather modest advantage; for example, for *N* = 10,000, the selective advantage is only 0.001. MR thus appears to be susceptible to being reversed by beneficial mutations of modest strength.

**Fig. 3.**
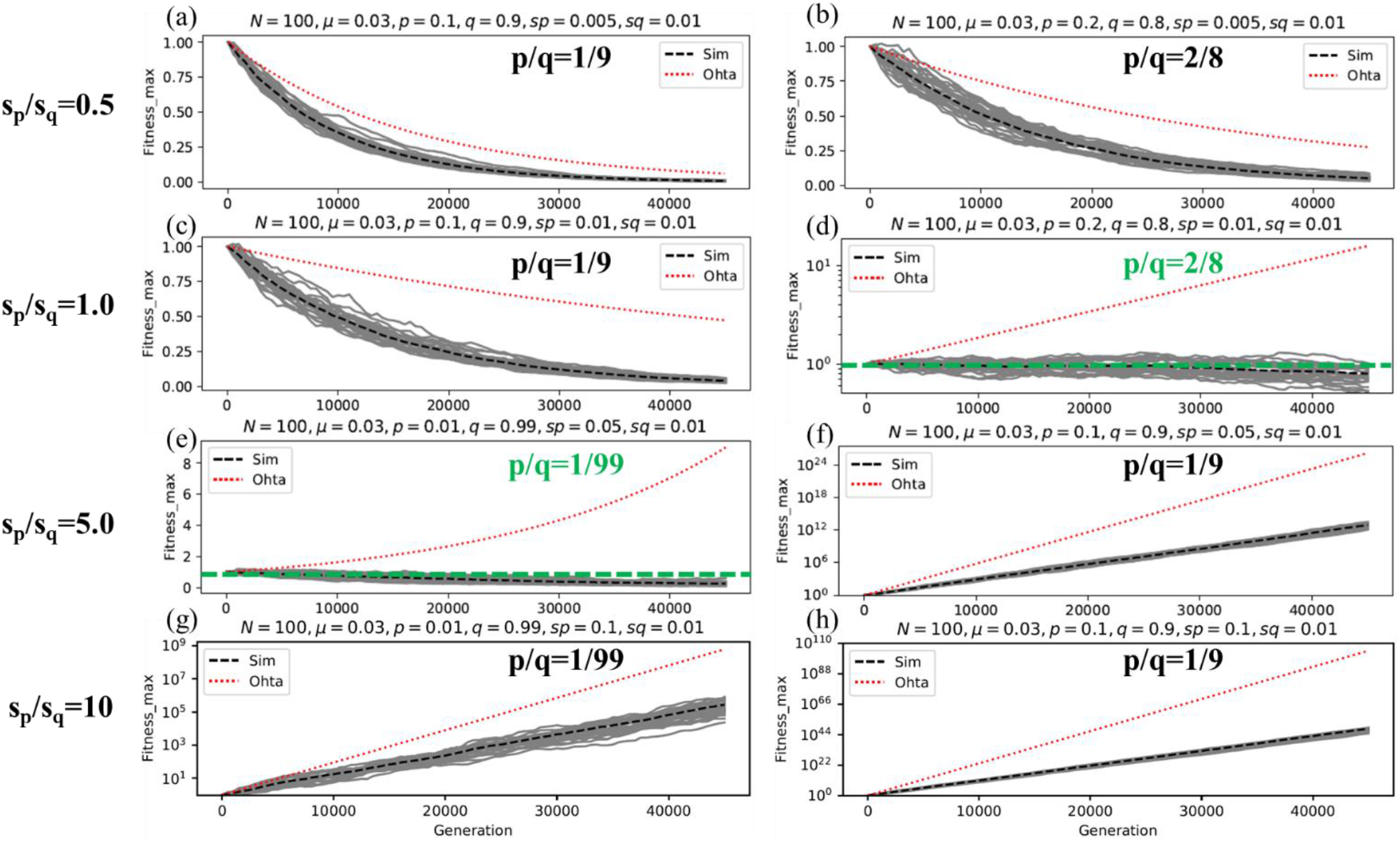
Fitness change with both beneficial and deleterious mutations. The red dashed line is the fitness dynamics of OR over time, obtained by Eq. (6’). The grey solid lines (25 repeats) are for MR, obtained by simulation (see Methods). In panels (a-c), the fitness would continue to decline for both MR and OR in spite of the input of beneficial mutations. In contrast (panels f-h), the fitness would rise by either process. Recombination plays a significant role only when MR is more robust than OR under the influence of beneficial mutations, as in panels (d-e) where the green dashed line shows the constant fitness at 1.

## Discussion

Since MR, as well as the more general HR process, is foremost about the power of recombination, a proper analysis should be the comparison of MR and OR, without and with recombination. The small degree of difference between the two processes seems intuitively obvious. As first shown by Haigh (1978), MR is effective when i) *N*, the population size, is small; ii) *s*, the selective strength, is weak; and iii) *U*, the genomic mutation rate, is large. These are also the conditions for OR to work effectively. Hence, it is not surprising that MR is biologically significant only in a small parameter space, vis-à-vis OR. In particular, when *Ns* < 1, OR is nearly equally effective. Since weakly deleterious mutations with *Ns* < 1 are rather common (Lu and Wu 2005; Eyre-Walker, et al. 2006; Eyre-Walker and Keightley 2007; Huber, et al. 2017), turning on full recombination is not likely to stall the ratchet, or the fitness decline.

In this study, we also consider the power of beneficial mutations in reversing the ratchet. While some authors may incorporate beneficial mutations into a more general MR, a ratchet is a devise of one-way movement. (MR with beneficial mutations is no longer a ratchet and should probably be called Muller’s Wrench.) In general, either ratchet can be easily reversed by beneficial mutations because beneficial mutation can impact the population fitness much more strongly than a deleterious mutation of equal strength in the opposite direction. The condition for reversal as specified in Eq. (7) is not particularly stringent. For that reason, the ratchet has to operate mostly between the emergence of beneficial mutations in order to avoid being reversed. In this study, the fitness effects of mutations, both beneficial and deleterious, are assumed to be multiplicative. While this assumption may not be realistic, the goal is to assess the difference in the efficacy of MR vs. OR. As suggested above, in the extreme form whereby beneficial mutations are epistatic over deleterious ones by the “bypass mechanism”, the difference may be even smaller than under the multiplicative model. Other forms of epistasis, such as the truncating selection model of Kondrashov (Kondrashov 1984; Kondrashov 1988), may also have similar impacts on MR and OR. In general, epistatic fitness interactions are expected to generate complex, and often unpredictable, outcomes (Chen, et al. 2019; He, et al. 2019).

In this analysis, we did not cover the range of mutation rate but set *u* close to that of Drosophila and human, respectively. Nevertheless, if *u* is very small, recombination would not matter, for instance, if there is only one polymorphic site in the genome. On the other hand, if *u* is large, then the chance of accruing beneficial mutations cannot be neglected. Eq. (7) only describes beneficial mutations of equal strength. However, if we assume the selective advantage to have a long-tailed distribution, then a large *u* would mean the availability of highly beneficial mutations to reverse the fitness decline.

In conclusion, the fitness decline generally associated with MR has much to do with the selection landscape, defined by the frequency and strength of fitness mutations, beneficial as well as deleterious. In a large part of this landscape, recombination is not a major factor. The significance of MR in the real world will therefore require detailed examinations of the various biological phenomena attributed to MR. A companion study covers the empirical evidence for MR (Wang et al. 2020; see Supplement).

## Methods

In this study, we simulate a discrete-time Wright-Fisher model for the MR effect as follows. At initial time, there are *N* individuals and every individual has no mutations. The fitness of an individual with *n*_*p*_ beneficial mutations and *n*_*q*_ deleterious mutations is 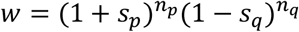, where and *s*_*p*_, *s*_*q*_ are the fitness gain and fitness loss of beneficial and deleterious mutations. Each generation consists of separate selection and mutation steps. At selection step, we will sample *N* individuals with replacement proportional to the relative fitness of each individual. And then each of the *N* individuals will reproduce one offspring to consist of the next generation. During the reproduction process, the mutation accumulation of each individual will follow Poisson distribution with rate *μ*. The proportions of beneficial mutations, deleterious, and neutral mutations will be *p*, *q*, and 1-*p*-*q*. Note *p*=0 represents the MR ratchet.

The above process is repeated (25 repeats for each parameter set unless otherwise specified) for a specified number of generations. Note it takes some time for the population to reach a steady state. Thus, we measure the features of dynamics after an equilibration time (10^4^ generations are sufficient in this study; see supplement) to remove transient effects from the initial conditions.

To estimate the accumulation rate of mutations, we will simulate 25 repeats for each parameter set and then calculate the average number of mutations of least-load-class individuals over time. The rate of mutation accumulation will be constant over time (black dashed lines in Fig. 2b and 2d). Having obtained the accumulation rate of mutations, we can calculate the fitness decline per generation for MR according to Eq. (8). Note that it’s much time-consuming while simulating the dynamics of a large population size (e.g. *N*=10000 or 100000). In this case, we only simulate 100000 generations. If the mutation number of least-load individual is still 0, we just estimate the rate of mutation accumulation to be less than 10^−5^.

## Acknowledgements

We thank Drs Jianzhi Zhang and Yunxin Fu for comments and suggestions on this manuscript. This work was funded by the National Natural Science Foundation of China (31730046, 91131000 to C.I.W., 31900417 to G.L., 81972691 to H.W.), Guangdong Basic and Applied Basic Research Foundation (2020B1515020030 to H.W., 2019A1515010708 to G.L.), National Key Research and Development Project (2020YFC0847000 to H.W.).

## References

Bachtrog D. 2013. Y-chromosome evolution: emerging insights into processes of Y-chromosome degeneration. Nature Reviews Genetics 14:113–124.

Charlesworth B, Betancourt AJ, Kaiser VB, Gordo I. 2009. Genetic recombination and molecular evolution. Cold Spring Harb Symp Quant Biol 74:177–186.

Charlesworth B, Charlesworth D. 2000. The degeneration of Y chromosomes. Philos Trans R Soc Lond B Biol Sci 355:1563–1572.

Chen Y, Shen Y, Lin P, Tong D, Zhao Y, Allesina S, Shen X, Wu C-I. 2019. Gene regulatory network stabilized by pervasive weak repressions: microRNA functions revealed by the May–Wigner theory. National Science Review 6:1176–1188.

Comeron JM, Williford A, Kliman RM. 2008. The Hill-Robertson effect: evolutionary consequences of weak selection and linkage in finite populations. Heredity (Edinb) 100:19–31.

Desai MM. 2020. Haigh (1978) and Muller’s ratchet. Theor Popul Biol 133:19–20.

Etheridge AM, Pfaffelhuber P, Wakolbinger A. 2009. How often does the ratchet click? Facts, heuristics, asymptotics. LONDON MATHEMATICAL SOCIETY LECTURE NOTE SERIES:365–390.

Eyre-Walker A, Keightley PD. 2007. The distribution of fitness effects of new mutations. Nat Rev Genet 8:610–618.

Eyre-Walker A, Woolfit M, Phelps T. 2006. The distribution of fitness effects of new deleterious amino acid mutations in humans. Genetics 173:891–900.

Felsenstein J. 1974. The evolutionary advantage of recombination. Genetics 78:737–756.

Gabriel W, Lynch M, Burger R. 1993. Muller’s Ratchet and Mutational Meltdowns. Evolution 47:1744–1757.

Gessler DD. 1995. The constraints of finite size in asexual populations and the rate of the ratchet. Genet Res 66:241–253.

Gillespie JH. 1998. Population Genetics: A Concise Guide: Johns Hopkins University Press.

Good BH, Walczak AM, Neher RA, Desai MM. 2014. Genetic diversity in the interference selection limit. PLoS Genet 10:e1004222.

Gordo I, Charlesworth B. 2000a. The degeneration of asexual haploid populations and the speed of Muller’s ratchet. Genetics 154:1379–1387.

Gordo I, Charlesworth B. 2000b. On the Speed of Muller’s Ratchet. Genetics 156:2137–2140.

Goyal S, Balick DJ, Jerison ER, Neher RA, Shraiman BI, Desai MM. 2012. Dynamic mutation-selection balance as an evolutionary attractor. Genetics 191:1309–1319.

Haag-Liautard C, Dorris M, Maside X, Macaskill S, Halligan DL, Charlesworth B, Keightley PD. 2007. Direct estimation of per nucleotide and genomic deleterious mutation rates in Drosophila. Nature 445:82–85.

Haigh J. 1978. The accumulation of deleterious genes in a population--Muller’s Ratchet. Theor Popul Biol 14:251–267.

He Z, Li X, Yang M, Wang X, Zhong C, Duke NC, Wu CI, Shi S. 2019. Speciation with gene flow via cycles of isolation and migration: insights from multiple mangrove taxa. Natl Sci Rev 6:275–288.

Hill WG, Robertson A. 1966. The effect of linkage on limits to artificial selection. Genet Res 8:269–294.

Huber CD, Kim BY, Marsden CD, Lohmueller KE. 2017. Determining the factors driving selective effects of new nonsynonymous mutations. Proc Natl Acad Sci U S A 114:4465–4470.

Jonsson H, Sulem P, Kehr B, Kristmundsdottir S, Zink F, Hjartarson E, Hardarson MT, Hjorleifsson KE, Eggertsson HP, Gudjonsson SA, et al. 2017. Parental influence on human germline de novo mutations in 1,548 trios from Iceland. Nature 549:519–522.

Keightley PD, Ness RW, Halligan DL, Haddrill PR. 2014. Estimation of the spontaneous mutation rate per nucleotide site in a Drosophila melanogaster full-sib family. Genetics 196:313–320.

Kondrashov AS. 1988. Deleterious mutations and the evolution of sexual reproduction. Nature 336:435–440.

Kondrashov AS. 1984. Deleterious mutations as an evolutionary factor: 1. The advantage of recombination. Genetical Research 44:199–217.

Leu J-Y, Chang S-L, Chao J-C, Woods LC, McDonald MJ. 2020. Sex alters molecular evolution in diploid experimental populations of S. cerevisiae. Nature Ecology & Evolution 4:453–460.

Lu J, Wu CI. 2005. Weak selection revealed by the whole-genome comparison of the X chromosome and autosomes of human and chimpanzee. Proc Natl Acad Sci U S A 102:4063–4067.

McDonald MJ, Rice DP, Desai MM. 2016. Sex speeds adaptation by altering the dynamics of molecular evolution. Nature 531:233–236.

McVean GAT, Charlesworth B. 2000. The effects of Hill-Robertson interference between weakly selected mutations on patterns of molecular evolution and variation. Genetics 155:929–944.

Muller HJ. 1964. The Relation of Recombination to Mutational Advance. Mutat Res 106:2–9.

Neher RA. 2013. Genetic Draft, Selective Interference, and Population Genetics of Rapid Adaptation. Annual Review of Ecology, Evolution, and Systematics, Vol 44 44:195–215.

Neher RA, Shraiman BI. 2012. Fluctuations of fitness distributions and the rate of Muller’s ratchet. Genetics 191:1283–1293.

Ohta T. 1992. The Nearly Neutral Theory of Molecular Evolution. Annual Review of Ecology and Systematics 23:263–286.

Ohta T. 1976. Role of very slightly deleterious mutations in molecular evolution and polymorphism. Theoretical Population Biology 10:254–275.

Ohta T. 1973. Slightly deleterious mutant substitutions in evolution. Nature 246:96–98.

Ohta T. 1987. Very slightly deleterious mutations and the molecular clock. Journal of Molecular Evolution 26:1–6.

Ruan Y, Wang H, Chen B, Wen H, Wu CI. 2020. Mutations Beget More Mutations-Rapid Evolution of Mutation Rate in Response to the Risk of Runaway Accumulation. Mol Biol Evol 37:1007–1019.

Scally A, Durbin R. 2012. Revising the human mutation rate: implications for understanding human evolution. Nat Rev Genet 13:745–753.

Shi L, Luo X, Jiang J, Chen Y, Liu C, Hu T, Li M, Lin Q, Li Y, Huang J, et al. 2019. Transgenic rhesus monkeys carrying the human MCPH1 gene copies show human-like neoteny of brain development. National Science Review 6:480–493.

Smith JM. 1978. The evolution of sex. London, Cambridge University Press.

Spielman SJ, Wilke CO. 2015. The relationship between dN/dS and scaled selection coefficients. Molecular Biology and Evolution 32:1097–1108.

Weissman DB, Hallatschek O. 2014. The rate of adaptation in large sexual populations with linear chromosomes. Genetics 196:1167–1183.

Wu CI, Wang HY, Ling S, Lu X. 2016. The Ecology and Evolution of Cancer: The Ultra-Microevolutionary Process. Annu Rev Genet 50:347–369.

Xu X, Li G, Li C, Zhang J, Wang Q, Simmons DK, Chen X, Wijesena N, Zhu W, Wang Z, et al. 2019. Evolutionary transition between invertebrates and vertebrates via methylation reprogramming in embryogenesis. National Science Review 6:993–1003.

