## Supplement for "Sex, fitness decline and recombination – Muller’s ratchet vs. Ohta’s ratchet"

### Supplementary Information includes:

Related Manuscript: Wang et al. (A separate but concurrent submission to MBE)

Supplementary text

Fig. S1 to S4

### Related Manuscript: Wang et al. (A separate but concurrent submission to MBE)

#### Muller's ratchet – Does it really operate in nature?

Haiyu Wang[1], Yongsen Ruan[1], Lingjie Zhang[1], Xuemei Lu[2,3], Haijun Wen[1*] and Chung-I Wu[1,4*]

[1]State Key Laboratory of Biocontrol, School of Life Sciences, Sun Yat-sen University, Guangzhou 510275, China.
[2]State Key Laboratory of Genetic Resources and Evolution, Kunming Institute of Zoology, Chinese Academy of Science, Kunming 650223, China.
[3]Center for Excellence in Animal Evolution and Genetics, Chinese Academy of Sciences, Kunming 650223, China.
[4]Department of Ecology and Evolution, University of Chicago, Chicago, Illinois 60637, USA

[*]Correspondence to:
Haijun Wen,
Chung-I Wu,

**Abstract** – Muller's ratchet (MR) refers to the irreversible decline of population fitness due to the accumulation of deleterious mutations in the absence of recombination. Since MR is in opposition to natural selection, it must operate within a very narrow parameter space. Outside of this space, either MR does not work or the ratchet mechanism can work well even with recombination (see Supplement). Therefore, MR may not be as powerful a driving force of evolution as believed. Here, we survey the empirical evidence for the key evolutionary events, including the emergence of sexual reproduction and the degeneracy of the Y chromosome, that have been attributed to the operation of MR. We found little empirical support for MR vis-à-vis other explanations. Overall, the impact of MR on major evolutionary events seems minimal. The one niche where MR might operate is the ultra-microevolution of somatic cells but the parameter space remains uncertain. In short, age-dependent fitness decline appears to be the last frontier for validating MR in nature.

### Introduction

H. J. Muller (1964) proposed a mechanism, now referred to as Muller's Ratchet (MR), that would allow deleterious mutations to accumulate, even though selection is working against the accumulation. In the conventional interpretation, the mechanism would work only in asexual populations as recombination in sexual species can reverse the accumulation among individuals (but see Ruan et al. 2020 in the Supplement). Under MR, the population fitness will continue to decline, thus necessitating genomic responses to reverse the trend.

This continual fitness decline makes MR significant for two reasons. First, the evolution in opposition to natural selection is of substantial theoretical interest (Charlesworth 1978; Haigh 1978; Gabriel, et al. 1993; Barton, et al. 1998; Loewe 2006), especially when MR is compared with a process with recombination (see the Supplement). It is intriguing whether the operating conditions for MR are realizable in real life. Second (and more important), many biological phenomena have been proposed to have evolved to avert MR and, hence, to avoid the fitness decline. These phenomena, including the evolution of sexual reproduction and the degeneracy of Y chromosome, are the main focus of this review.

MR is the subject of a substantial literature, which has shown the working of MR in laboratory settings where the parameter values, such as the population size ($N$), are rather extreme (Mukai, et al. 1972; Felsenstein 1974; Haigh 1978; Kondrashov 1988; Chao 1990; Lynch, et al. 1993; Lynch, et al. 1995; Andersson, et al. 1996; Keightley, et al. 1997; Zeyl, et al. 2001; Bank, et al. 2016; Zhang, et al. 2019). Obviously, MR should work if the population goes through bottlenecks of $N = 1$ continually. It is far less certain whether MR works in nature where the parameter values are rarely so extreme. In this review, we re-assess the significance of MR in evolutionary biology.

### The concept and operation of MR

#### *What is (or is not) MR?*

The essence of MR is a continual and irreversible decline in the fitness of the population. The mechanism that drives the decline is the accumulation of deleterious mutations in the absence of recombination (Muller 1964; Felsenstein 1974; Haigh 1978; Gabriel, et al. 1993; Jain 2008).

Since MR is expected to lead to the decline of population fitness, many phenomena attributed to MR may not qualify. For example, the accumulation of recessive mutations in asexual diploids with little fitness decline should not be considered an example of MR (Zeyl, et al. 2001; Lovell, et al. 2017). The requisite fitness decline points to the main difficulty in demonstrating MR's biological significance: any phenomenon posited to be driven by MR must have successfully evaded its action. In other words, one expects to identify MR's impact by not finding MR in operation. Indeed, Muller (1964) may be the first to suggest that sexual reproduction and recombination may have evolved to suppress MR.

MR is related to the more general process of mutation interferences, often referred to as the Hill-Robertson effect (HR, (Felsenstein 1974; McVean, et al. 2000; Lu, et al. 2006; Charlesworth, et al. 2009)). Charlesworth et al. (2009) suggested the interferences among beneficial and deleterious mutations as the related processes (their Table 1). MR is nevertheless a distinct concept, the essence of which being the continual fitness decline as well as the biological phenomena associated with this hypothetical decline.

*How does MR work?*

MR operates mainly in very small populations. Its efficacy decreases in large populations as summarized by a simple equation (Haigh 1978)

$$n_{min} = N \exp(-U/s) \qquad (1)$$

where $N$ is the population size, $U$ is the rate of deleterious mutation and $s$ is the selective intensity against the mutation. In Eq. (1), $n_{min}$ is the number of individuals in the population with the smallest number (min) of mutations. This least loaded class represents the fittest individuals in the population. When $n_{min}$ becomes 0, $n_{min+1}$ becomes the new $n_{min}$ and the ratchet moves a notch. The farther the ratchet moves, the lower the fitness of the population becomes. Clearly, a very small $n_{min}$ is more likely to become 0 and, hence, the operation of MR depends on $N$, $U$ and $s$. It has been pointed out that the movement of the ratchet does not necessarily mean the fixation of deleterious mutations (Charlesworth, et al. 1992). By Eq. (1), the conditions for MR to work are i) $N$ and $s$ being sufficiently small, and/or ii) $U$ being sufficiently large.

### The conceptual and operational limitations of MR

*1) Ohta' ratchet – The continual fitness decline with free recombination*

While the central tenet of MR is the absence of recombination facilitating the decline of population fitness, a companion study (Ruan et al. 2020; Appended in the beginning of the Supplement) asks the obvious question: If the population suddenly gains the ability to recombine freely, does the fitness decline stop? In the full parameter space of population size, selection and mutation, Ruan et al. found that a freely recombining population may often experience the irreversible fitness decline generally attributed to MR. The process is referred to as Ohta's ratchet (OR) in honor of Tomoko Ohta's contributions to the study of slightly deleterious mutations (Ohta 1973, 1976, 1987, 1992). MR operates more effectively than OR only in a limited parameter space. In a realistic evolutionary process, the fitness decline may not be stopped by gaining the capacity to recombine.

*2) MR in relation to beneficial mutations*

Although MR is presented as a cumulative process of deleterious mutations, the same mutational process would certainly generate beneficial mutations as well. One approach is to expand the model to incorporate beneficial mutations (Goyal, et al. 2012). However, the expansion would violate the central tenet of the MR process since a ratchet is first and foremost an irreversible mechanism. Instead, Muller (1964) and Haigh (1978) treat beneficial mutations as external forces that oppose MR, a practice adopted in this study. The semantic difference between the two approaches would not affect the answer to the key question posed below.

How effectively can beneficial mutations stall the ratchet? The answer would depend on the frequency and strength of beneficial mutations. In general, a beneficial mutation has an overwhelming effect over a deleterious one that changes the fitness by the same amount but in the opposite direction. For example, if the fitness gain/loss is 0.001 and the population size is 10,000, then a beneficial mutation would require ~ $2 \times 10^9$ deleterious mutations to offset the gain. In a wider parameter space, Ruan et al. (2020; see Supplement) show that beneficial mutations can usually stop and reverse the ratchets. For that reason, the parameter space where MR works, but OR does not, is important and the space is restricted (see Ruan et al. in Supplement for detail.)

*3) Empirical observations on beneficial mutations*

As MR is driven by deleterious mutations and opposed by beneficial mutations, it must operate before the population accrues beneficial mutations of sufficient strength. (Weak beneficial mutations can be treated as neutral mutations in the context of MR.) Obviously, this “waiting time” between beneficial mutations must be long enough for biological phenomena (such as sexual reproduction) to evolve in response to the fitness decline.

While beneficial mutations used to be viewed as being extremely rare, they appear rather common in laboratory experiments on *E. coli* (Barrick, et al. 2010; Sprouffske, et al. 2018) . Recent genomic data have also led to a revision of this view (Fay, et al. 2001; Fay, et al. 2002; Eyre-Walker, et al. 2007; Schneider, et al. 2011; Enard, et al. 2014). For example, 30% of fixed nonsynonymous mutations have been estimated to be beneficial in *Drosophila* (Fay, et al. 2002). It is also plausible that beneficial mutations might be even more common when the population is sagged with deleterious mutations (Peck 1994; Joseph, et al. 2004).

A critical issue is the interactions between beneficial and deleterious mutations, which are generally assumed to be multiplicative. Multiplicative fitness would be reasonable if mutations effects are weak, i.e., small-step changes (Fisher 1930; Kimura 1983; Wagner, et al. 2011; He, et al. 2019; Xu, et al. 2019). Recently, using the physico-chemical distances between amino acids as the step sizes of evolution, Chen et al. (Chen, He, et al. 2019; Chen, Lan, et al. 2019) have reached a different conclusion. While negative selection indeed tolerates small step changes across a wide spectrum of taxa (Chen, Lan, et al. 2019), positive selection does not favor small-step mutations (Chen, He, et al. 2019). The findings may support the use of a multiplicative fitness scale on deleterious mutations, but the scale should not be extended to beneficial mutations.

In mechanistic terms, beneficial mutations may simply “by-pass” the deleterious effects, thus invalidating the multiplicative model. For example, if deleterious effects happen in one environment but the beneficial mutations cause the organisms to move to a different environment, then the two types of fitness effect are not multiplicative. It may not be unreasonable to view some beneficial mutations to reset the fitness landscape as if to restart the adaptive evolution in new environments. In domesticated species (Lu, et al. 2006), deleterious mutations hitchhike with the beneficial ones, rather than drag them down. In the evolution of Y chromosomes, beneficial mutations that are male biased appear to operate separately from the deleterious ones (Bachtrog 2013).

The interactions between beneficial and deleterious mutations can be most readily analyzed in the evolution of somatic cells. Across more than 10,000 tumor samples from the TCGA database (Cancer Genome Atlas 2012; Cancer Genome Atlas Research, et al. 2013; Kandoth, et al. 2013; Lawrence, et al. 2014), the number of nonsynonymous mutations ranges between < 30 to > 3000. The proportion of non-neutral mutations in the coding regions is ~ 20% (Wu, et al. 2016) and the ratio of beneficial : deleterious mutations has been estimated to be 1 : (4-5) (Wu, et al. 2016; Chen, Shi, et al. 2019). Hence, for the more highly loaded cases, the tumors would have about ~ 500 deleterious and ~ 100 beneficial mutations. Given such a large number of fitness-altering mutations, it would seem implausible that the fitness effects would be multiplicative.

In this perspective, beneficial mutations are those that can elevate the recipients' fitness above the most-fit class in the population and may continue to improve the population fitness in the presence of (previously) deleterious mutations. By this definition, MR could operate only between the emergences of beneficial mutations.

**Re-evaluation of the empirical evidence for phenomena attributed to MR**

There are six biological phenomena attributed to MR which we shall review below.

1. The evolution of sex and recombination

MR was initially proposed to explain sexual reproduction and the concomitant genetic recombination. Muller suggested that sex and recombination can reverse the movement of the ratchet. Subsequent theoretical developments have shown that recombination under some conditions can be so beneficial that it would compensate for the inherent "two-fold" cost of sexual reproduction (Maynard Smith 1978; Gillespie 2004).

Nevertheless, the cause and consequence may have been reversed in this narrative. In other words, sex and recombination may have evolved for other reasons with the annulment of MR being a consequence. Indeed, there are many explanations for the evolution of sex (Maynard Smith 1978); for example, the diversification of genotypes facing environmental changes. These other explanations are beyond the scope of this survey.

A more realistic application of the MR model should be the maintenance, rather than the origin, of sexual reproduction. This more modest proposition is testable by surveying species with facultative sexual reproduction. The switch from asexual to sexual reproduction in these species does not require the evolution of sexual reproduction from scratch; a phenotypic switch would suffice. Testing whether this phenotypic switch is a response to the movement of MR should shed light on MR and sexual reproduction. Thirty years ago, G. Bell (1989) famously supported this hypothesis by presenting extensive observations on the switching of reproductive modes in protozoans. Although his interpretations were not enthusiastically accepted, it was hoped that "this book will stimulate further experiments" on the hypothesis (Lenski 1990). We now revisit the topic.

2. The phenotypic switch between sexual and asexual reproduction

Many facultative sexual species predominantly adopt the asexual mode and switch to sexual reproduction only at intervals (Hadany, et al. 2007; Cáceres, et al. 2009). This switch has often been interpreted to be a means of reversing the fitness decline, when the deleterious mutation load approaches a critical level. Such a critical level is an essential feature in, for example, Kondrashov's (1988) model of transition from asexual to sexual reproduction. Similarly, in asexual (apomictic) plants, a small fraction of sexually reproducing individuals can often be found amidst the majority of asexually reproducing ones (Wright, et al. 2013; Hojsgaard, et al. 2015).

However, the empirical evidence is generally incompatible with the hypothesis that facultative species switch on sexual reproduction to reverse MR. If MR is the cause of the switch, one would have expected the switch to happen predominantly in small populations that grow in the same environment for a certain length of time. However, the general pattern is that populations would make the switch when facing new environmental stresses, regardless of the population size. This is true in protozoans (Bell 1989; Carter, et al. 2013; Orias, et al. 2017), rotifers (Snell, et al. 2006; Becks, et al. 2010; Becks, et al. 2012; Luijckx, et al. 2017), yeast (Bernstein, et al. 1989; Neiman 2011) and apomictic plants (Pellino, et al. 2013; Bishop, et al. 2017). In other words, the switch appears to happen in the wrong populations and wrong environments, according to the MR model.

Overall, the switch to sexual reproduction has been more convincingly demonstrated to be associated with the diversification of genotypes such that some of the new ones would be able to adapt to the new environment (Wojciechowski, et al. 1989; Kleunen, et al. 2001; Jarmer, et al. 2002; Grishkan, et al. 2003; Foster 2005; Goddard, et al. 2005; King, et al. 2009; Berman, et al. 2012).

3. The permanent return to asexual reproduction

According to the MR model, it is unlikely that a sexual species would return to asexual reproduction permanently. Hence, cases where the permanent return has been recorded will be informative about the mechanisms by which MR is rendered non-operative.

A well-known case is the bdelloid rotifers, which has returned to asexuality that has lasted long enough to be considered permanent (Flot, et al. 2013; Schwander 2016). There have been many adaptive features in the rotifer genomes, suggesting that beneficial mutations have been common in their return to asexuality. A direct proof of the frequent occurrences of beneficial mutations can be found in yeast populations as asexual lines often end up with a higher population fitness than in the beginning of the experiment (Zeyl, et al. 2001).

Among protozoans, many strains from several Tetrahymena species have lost the micro-nuclei, which are necessary for sexual reproduction (Doerder 2014; Orias, et al. 2017). The divergence in the macro-nuclei also suggests the long-term return to asexuality (Doerder 2014; Orias, et al. 2017), much like the rotifers. Asexual strains of Tetrahymena, however, use a peculiar mechanism of amitosis by which chromosomes in the high-ploidy macro-nuclei are randomly assorted. When cells divide, such assortments can yield a “less loaded genotype” than the parent (Doerder 2014). In other words, the asexual strains of Tetrahymena simply find a way to reverse the ratchet, further suggesting the limitations on MR’s operation.

4. MR in selfing species

With proper scaling of the parameters and under conditions more stringent than in asexual species, MR can also work in selfing taxa (Heller, et al. 1979; Charlesworth, et al. 1992). In the MR model, deleterious mutations would accumulate in the selfing phase. As the ratchet progresses, outcrossing would be beneficial, as it reduces the mutation load. The switching from selfing to outcrossing has been observed regularly but rarely has this switch been attributed to the MR mechanism. This is true in nematodes (Cutter 2005; Morran, et al. 2009; Morran, et al. 2011) and plants (Bishop, et al. 2016; Bishop, et al. 2017).

5. The degeneracy of Y chromosome

The degeneracy of Y chromosome has happened independently multiple times (Charlesworth 1996; Charlesworth, et al. 2000; Bachtrog 2013) and is still happening in the evolution of the neo-Y chromosome (Bachtrog 2006; Zhou, et al. 2012; Bachtrog 2013; Hough, et al. 2014). MR would be an obvious explanation for the Y-degeneracy but the reservations about MR discussed above could apply to Y degeneracy as well. Indeed, the observation on polymorphism does not support the operation of MR as the reduced Y-linked polymorphism is attributed to either background selection (Hough et al, 2017) or selective sweep.

Among the many competing hypotheses, one is the hitchhiking hypothesis whereby a beneficial mutation on the Y may drag along degenerated loci to fixation. In this scenario, deleterious mutations would be common on the Y chromosome as widely reported. However, the fixation is not driven by MR but by hitchhiking with linked beneficial mutations (Bachtrog 2013). In fact, if MR is the mechanism, Y degeneracy should be observed mainly in species with a small population size but Y degeneracy seems too common to fit this explanation. Hence, there is no compelling evidence that MR is the best explanation for this fairly common phenomenon of Y degeneracy (Charlesworth, et al. 2000; Kaiser, et al. 2009; Bachtrog 2013; Hough, et al. 2017).

6. Genomic features

It has also been suggested that protozoans, bacteria, plants and even cancer cells develop polyploidy to mitigate against the “ravage” of deleterious mutations (Maciver 2016; López, et al. 2019), presumably on accout of transcriptome stability (Chen, Shen, et al. 2019). In this interpretation, polyploidy evolves first and a consequence is that MR would be stalled. Thus, MR is not a driving force of evolution in this context.

**The last frontier for MR - Aging and somatic cell evolution**

As stated in Introduction, the potential impact of MR in biology rests on the ability of biological systems to evade its action. This partially explains why it has been so difficult to prove the causative link between MR and sex/recombination. This dilemma can be resolved if the phenomenon hypothesized to be driven by MR is associated with an ongoing fitness decline. In this perspective, we suggest that the evolution of cells within a multi-cellular organism may be a natural process driven by MR. Somatic cell evolution follows rules that are similar, but not identical, to those governing organismal evolution. The evolution is ultra-microevolutionary in scale (Wu, et al. 2016) and its characteristics germane to MR are as follows.

First, there is indeed a fitness decline associated with aging in many parts of somatic tissues (Henry, et al. 2010; Henry, et al. 2015; Cannataro, et al. 2016). For example, the bone marrow in older mice has a reduced output of B-progenitor cells (Henry et al. 2010, 2015), suggesting deleterious changes in the stem cell niches.

Second, the unit of somatic evolution appears to be the stem cell niche (Cairns 1975; Greaves 2013) and the size of the stem cell population ($N$) is small. In the colon, $N < 50$ (Kozar, et al. 2013; Chen, Shi, et al. 2019) which would be small enough to allow MR to move quickly. At this range of $N$, selection efficacy is very low and the evolutionary rate is very close to the neutral rare, referred to as quasi-neutrality (Chen, Shi, et al. 2019). In the regime of weak selection, MR operates quite effectively. For a side note, $N$ in the stem cell niche can grow from $< 10^2$ to $> 10^9$ if the population evolves to become tumor and MR would cease to work (Ling, et al. 2015; Wang, et al. 2017; Wen, et al. 2018). In this sense, MR holds tumorigenesis in check at the early stage but not after the tumor evolves “out of the gate”.

Third, there are millions of stem cell niches on the epithelial of the colon alone. For that reason, events with a very low probability per small population may not be rare in aggregate. The large number of evolutionary units experiencing fitness decline will make detection much more feasible.

Fourth, the mutation rate in the soma is typically 100 fold higher than the germline mutation rate (Cancer Genome Atlas 2012; Blokzijl, et al. 2016; Ruan, et al. 2020). Note that small $N$ and large $U$ are the ideal conditions for the operation of MR. Fifth, somatic cell evolution happens at the ultra-microevolutionary scale of one life span. In this duration, some of the stem cell populations might evolve according to the rules of MR without being interrupted by beneficial mutations.

**Concluding remarks**

In the five decades since the idea of MR was proposed, the topic has been heavy in theory but relatively slim on the empirical side. In the recent literature, MR has increasingly been associated with somatic cell evolution (Cannataro, et al. 2016; Wu, et al. 2016; Zhang, et al. 2019). We have recently measured the fitness components of MR in the HeLa cell line (Zhang, et al. 2019) and further demonstrate the working of MR in cell populations of a moderate size (Wang HY et al. in preparation). Nevertheless, actual aging demands rapid ratchet movements in vivo in a lifetime and this speedy movement remains to be investigated. In this sense, aging of somatic tissue appears to be the last frontier where MR may be biologically relevant.

**Acknowledgments**

We would like to thank the members of Wu Lab for discussions and advices. This work was supported by the National Natural Science Foundation of China (31730046 and 91731301) and the 985 Project (33000-18841204). The

Key Research Program of the Chinese Academy of Sciences grant (KFZD-SW-220-1), National Natural Science Foundation of China grants (31771416) and CAS "Light of West China" Program.

**References**

Andersson DI, Hughes D. 1996. Muller's ratchet decreases fitness of a DNA-based microbe. Proceedings of the National Academy of Sciences 93:906-907.

Bachtrog D. 2006. Expression profile of a degenerating neo-Y chromosome in *Drosophila*. Curr Biol 16:1694-1699.

Bachtrog D. 2013. Y-chromosome evolution: emerging insights into processes of Y-chromosome degeneration. Nat Rev Genet 14:113-124.

Bank C, Renzette N, Liu P, Matuszewski S, Shim H, Foll M, Bolon DN, Zeldovich KB, Kowalik TF, Finberg RW, et al. 2016. An experimental evaluation of drug-induced mutational meltdown as an antiviral treatment strategy. Evolution 70:2470-2484.

Barrick JE, Kauth MR, Strelioff CC, Lenski RE. 2010. *Escherichia coli* rpoB mutants have increased evolvability in proportion to their fitness defects. Mol Biol Evol 27:1338-1347.

Barton NH, Charlesworth B. 1998. Why sex and recombination? Science 281:1986.

Becks L, Agrawal AF. 2012. The evolution of sex is favouredduring adaptation to new environments. PLOS Biology 10:e1001317.

Becks L, Agrawal AF. 2010. Higher rates of sex evolve in spatially heterogeneous environments. Nature 468:89-92.

Bell G. 1989. Sex and death in protozoa:the history of an obsession. Cambridge: Cambridge University Press.

Berman J, Hadany L. 2012. Does stress induce (para)sex? Implications for *Candida albicans* evolution. Trends Genet 28:197-203.

Bernstein C, Johns V. 1989. Sexual reproduction as a response to H2O2 damage in *Schizosaccharomyces* pombe. Journal of Bacteriology 171:1893.

Bishop J, Jones HE, Lukac M, Potts SG. 2016. Insect pollination reduces yield loss following heat stress in faba bean (*Vicia faba L*.). Agric Ecosyst Environ 220:89-96.

Bishop J, Jones HE, O'Sullivan DM, Potts SG. 2017. Elevated temperature drives a shift from selfing to outcrossing in the insect-pollinated legume, faba bean (*Vicia faba*). J Exp Bot 68:2055-2063.

Blokzijl F, de Ligt J, Jager M, Sasselli V, Roerink S, Sasaki N, Huch M, Boymans S, Kuijk E, Prins P, et al. 2016. Tissue-specific mutation accumulation in human adult stem cells during life. Nature 538:260-264.

Cáceres CE, Hartway C, Paczolt KA. 2009. Inbreeding depression varies with investment in sex in a facultative parthenogen. Evolution 63:2474-2480, 2477.

Cairns J. 1975. Mutation selection and the natural history of cancer. Nature 255:197-200.

Cancer Genome Atlas N. 2012. Comprehensive molecular characterization of human colon and rectal cancer. Nature 487:330-337.

Cancer Genome Atlas Research N, Weinstein JN, Collisson EA, Mills GB, Shaw KR, Ozenberger BA, Ellrott K, Shmulevich I, Sander C, Stuart JM. 2013. The Cancer Genome Atlas Pan-Cancer analysis project. Nat Genet 45:1113-1120.

Cannataro VL, McKinley SA, St Mary CM. 2016. The implications of small stem cell niche sizes and the distribution of fitness effects of new mutations in aging and tumorigenesis. Evol Appl 9:565-582.

Carter LM, Kafsack BF, Llinas M, Mideo N, Pollitt LC, Reece SE. 2013. Stress and sex in malaria parasites: Why does commitment vary? Evol Med Public Health 2013:135-147.

Chao L. 1990. Fitness of RNA virus decreased by Muller's ratchet. Nature 348:454-455.

Charlesworth B. 1996. The evolution of chromosomal sex determination and dosage compensation. Current Biology 6:149-162.

Charlesworth B. 1978. Model for evolution of Y chromosomes and dosage compensation. Proceedings of the National Academy of Sciences of the United States of America 75:5618-5622.

Charlesworth B, Betancourt AJ, Kaiser VB, Gordo I. 2009. Genetic recombination and molecular evolution. Cold Spring Harb Symp Quant Biol 74:177-186.

Charlesworth B, Charlesworth D. 2000. The degeneration of Y chromosomes. Philos Trans R Soc Lond B Biol Sci 355:1563-1572.

Charlesworth D, Morgan MT, Charlesworth B. 1992. The effect of linkage and population size on inbreeding depression due to mutational load. Genetical Research 59:49-61.

Chen B, Shi Z, Chen Q, Shen X, Shibata D, Wen H, Wu CI. 2019. Tumorigenesis as the paradigm of quasi-neutral molecular evolution. Mol Biol Evol 36:1430-1441.

Chen Q, He Z, Lan A, Shen X, Wen H, Wu CI. 2019. Molecular evolution in large steps-codon substitutions under positive selection. Mol Biol Evol 36:1862-1873.

Chen Q, Lan A, Shen X, Wu CI. 2019. Molecular evolution in small steps under prevailing negative selection: a nearly universal rule of codon substitution. Genome Biol Evol 11:2702-2712.

Chen Y, Shen Y, Lin P, Tong D, Zhao Y, Allesina S, Shen X, Wu C-I. 2019. Gene regulatory network stabilized by pervasive weak repressions: microRNA functions revealed by the May–Wigner theory. National Science Review 6:1176-1188.

Cutter AD. 2005. Mutation and the experimental evolution of outcrossing in *Caenorhabditis elegans*. J Evol Biol 18:27-34.

Doerder FP. 2014. Abandoning sex: multiple origins of asexuality in the ciliate *Tetrahymena*. BMC Evol Biol 14:112.

Enard D, Messer PW, Petrov DA. 2014. Genome-wide signals of positive selection in human evolution. Genome Res 24:885-895.

Eyre-Walker A, Keightley PD. 2007. The distribution of fitness effects of new mutations. Nat Rev Genet 8:610-618.

Fay JC, Wyckoff GJ, Wu C-I. 2002. Testing the neutral theory of molecular evolution with genomic data from *Drosophila*. Nature 415:1024-1026.

Fay JC, Wyckoff GJ, Wu CI. 2001. Positive and negative selection on the human genome. Genetics 158:1227-1234.

Felsenstein J. 1974. The evolutionary advantage of recombination. Genetics 78:737-756.

Fisher RAS. 1930. The genetical theory of natural selection. Oxford: Clarendon Press.

Flot JF, Hespeels B, Li X, Noel B, Arkhipova I, Danchin EG, Hejnol A, Henrissat B, Koszul R, Aury JM, et al. 2013. Genomic evidence for ameiotic evolution in the bdelloid rotifer *Adineta vaga*. Nature 500:453-457.

Foster PL. 2005. Stress responses and genetic variation in bacteria. Mutat Res 569:3-11.

Gabriel W, Lynch M, Bürger R. 1993. Muller's ratchet and mutational meltdowns. Evolution 47:1744-1757.

Gillespie JH. 2004. Population genetics: a concise guide second edition. Baltimore: Johns Hopkins University Press.

Goddard MR, Godfray HCJ, Burt A. 2005. Sex increases the efficacy of natural selection in experimental yeast populations. Nature 434:636-640.

Goyal S, Balick DJ, Jerison ER, Neher RA, Shraiman BI, Desai MM. 2012. Dynamic mutation-selection balance as an evolutionary attractor. Genetics 191:1309-1319.

Greaves M. 2013. Cancer stem cells as 'units of selection'. Evol Appl 6:102-108.

Grishkan I, Korol AB, Nevo E, Wasser SP. 2003. Ecological stress and sex evolution in soil microfungi. Proc Biol Sci 270:13-18.

Hadany L, Otto SP. 2007. The evolution of condition-dependent sex in the face of high costs. Genetics 176:1713-1727.

Haigh J. 1978. The accumulation of deleterious genes in a population—Muller's ratchet. Theoretical Population Biology 14:251-267.

He Z, Li X, Yang M, Wang X, Zhong C, Duke NC, Wu CI, Shi S. 2019. Speciation with gene flow via cycles of isolation and migration: insights from multiple mangrove taxa. National Science Review 6:275-288.

Heller R, Smith JM. 1979. Does Muller's ratchet work with selfing? Genetical Research 32:289-293.

Henry CJ, Casás-Selves M, Kim J, Zaberezhnyy V, Aghili L, Daniel AE, Jimenez L, Azam T, McNamee EN, Clambey ET, et al. 2015. Aging-associated inflammation promotes selection for adaptive oncogenic events in B cell progenitors. Journal of Clinical Investigation 125:4666-4680.

Henry CJ, Marusyk A, Zaberezhnyy V, Adane B, DeGregori J. 2010. Declining lymphoid progenitor fitness promotes aging-associated leukemogenesis. Proceedings of the National Academy of Sciences of the United States of America 107:21713-21718.

Hojsgaard D, Hörandl E. 2015. A little bit of sex matters for genome evolution in asexual plants. Frontiers in plant science 6:82-82.

Hough J, Hollister JD, Wang W, Barrett SC, Wright SI. 2014. Genetic degeneration of old and young Y chromosomes in the flowering plant *Rumex hastatulus*. Proceedings of the National Academy of Sciences 111:7713-7718.

Hough J, Wang W, Barrett SCH, Wright SI. 2017. Hill-Robertson Interference reduces genetic diversity on a young plant Y-chromosome. Genetics 207:685-695.

Jain K. 2008. Loss of least-loaded class in asexual populations due to drift and epistasis. Genetics 179:2125-2134.

Jarmer H, Berka R, Knudsen S, Saxild HH. 2002. Transcriptome analysis documents induced competence of Bacillus subtilis during nitrogen limiting conditions. FEMS Microbiology Letters 206:197-200.

Joseph SB, Hall DW. 2004. Spontaneous mutations in diploid *Saccharomyces cerevisiae*: more beneficial than expected. Genetics 168:1817-1825.

Kaiser VB, Charlesworth B. 2009. The effects of deleterious mutations on evolution in non-recombining genomes. Trends Genet 25:9-12.

Kandoth C, McLellan MD, Vandin F, Ye K, Niu B, Lu C, Xie M, Zhang Q, McMichael JF, Wyczalkowski MA, et al. 2013. Mutational landscape and significance across 12 major cancer types. Nature 502:333-339.

Keightley PD, Caballero A. 1997. Genomic mutation rates for lifetime reproductive output and lifespan in Caenorhabditis elegans. Proceedings of the National Academy of Sciences 94:3823.

Kimura M. 1983. The neutral theory of molecular evolution. Cambridge: Cambridge University Press.

King KC, Delph LF, Jokela J, Lively CM. 2009. The geographic mosaic of sex and the Red Queen. Curr Biol 19:1438-1441.

Kleunen Mv, Fischer M, Schmid B. 2001. Effects of intraspecific competition on size variation and reproductive allocation in a clonal plant. Oikos 94:515-524.

Kondrashov AS. 1988. Deleterious mutations and the evolution of sexual reproduction. Nature 336:435-440.

Kozar S, Morrissey E, Nicholson AM, van der Heijden M, Zecchini HI, Kemp R, Tavare S, Vermeulen L, Winton DJ. 2013. Continuous clonal labeling reveals small numbers of functional stem cells in intestinal crypts and adenomas. Cell Stem Cell 13:626-633.

López S, Lim E, Huebner A, Dietzen M, Mourikis T, Watkins TBK, Rowan A, Dewhurst SM, Birkbak NJ, Wilson GA, et al. 2019. Whole genome doubling mitigates Muller's ratchet in cancer evolution. bioRxiv:513457.

Lawrence MS, Stojanov P, Mermel CH, Robinson JT, Garraway LA, Golub TR, Meyerson M, Gabriel SB, Lander ES, Getz G. 2014. Discovery and saturation analysis of cancer genes across 21 tumour types. Nature 505:495-501.

Lenski RE. 1990. Sex and death in protozoa. The history of an obsession. Graham Bell. Cambridge University Press, New York, 1989. xiv, 199 pp., illus. Science 248:901.

Ling S, Hu Z, Yang Z, Yang F, Li Y, Lin P, Chen K, Dong L, Cao L, Tao Y, et al. 2015. Extremely high genetic diversity in a single tumor points to prevalence of non-Darwinian cell evolution. Proceedings of the National Academy of Sciences 112:E6496.

Loewe L. 2006. Quantifying the genomic decay paradox due to Muller's ratchet in human mitochondrial DNA. Genetical Research 87:133-159.

Lovell JT, Williamson RJ, Wright SI, McKay JK, Sharbel TF. 2017. Mutation accumulation in an asexual relative of *Arabidopsis*. PLoS Genet 13:e1006550.

Lu J, Tang T, Tang H, Huang J, Shi S, Wu CI. 2006. The accumulation of deleterious mutations in rice genomes: a hypothesis on the cost of domestication. Trends Genet 22:126-131.

Luijckx P, Ho EKH, Gasim M, Chen S, Stanic A, Yanchus C, Kim YS, Agrawal AF. 2017. Higher rates of sex evolve during adaptation to more complex environments. Proceedings of the National Academy of Sciences 114:534.

Lynch M, Bürger R, Butcher D, Gabriel W. 1993. The mutational meltdown in asexual populations. Journal of Heredity 84:339-344.

Lynch M, Conery J, Burger R. 1995. Mutational meltdowns in sexual populations. Evolution 49:1067-1080.

Maciver SK. 2016. Asexual amoebae escape Muller's ratchet through polyploidy. Trends Parasitol 32:855-862.

Maynard Smith J. 1978. The Evolution of Sex. Cambridge: Cambridge University Press.

McVean GAT, Charlesworth B. 2000. The effects of Hill-Robertson Interference between weakly welected mutations on patterns of molecular evolution and variation. Genetics 155:929.

Morran LT, Cappy BJ, Anderson JL, Phillips PC. 2009. Sexual partners for the stressed: facultative outcrossing in the self-fertilizing nematode *Caenorhabditis elegans*. Evolution 63:1473-1482.

Morran LT, Schmidt OG, Gelarden IA, Parrish RC, 2nd, Lively CM. 2011. Running with the Red Queen: host-parasite coevolution selects for biparental sex. Science 333:216-218.

Mukai T, Chigusa SI, Mettler LE, Crow JF. 1972. Mutation rate and dominance of genes affecting viability in *Drosophila melanogaster*. Genetics 72:335.

Muller HJ. 1964. The relation of recombination to mutational advance. Mutation Research/Fundamental and Molecular Mechanisms of Mutagenesis 1:2-9.

Neiman AM. 2011. Sporulation in the budding yeast *Saccharomyces cerevisiae*. Genetics 189:737.

Ohta T. 1992. The nearly neutral theory of molecular evolution. Annual Review of Ecology and Systematics 23:263-286.

Ohta T. 1976. Role of very slightly deleterious mutations in molecular evolution and polymorphism. Theoretical Population Biology 10:254-275.

Ohta T. 1973. Slightly deleterious mutant substitutions in evolution. Nature 246:96-98.

Ohta T. 1987. Very slightly deleterious mutations and the molecular clock. Journal of Molecular Evolution 26:1-6.

Orias E, Singh DP, Meyer E. 2017. Genetics and epigenetics of mating type determination in *Paramecium* and *Tetrahymena*. Annu Rev Microbiol 71:133-156.

Peck JR. 1994. A ruby in the rubbish: beneficial mutations, deleterious mutations and the evolution of sex. Genetics 137:597.

Pellino M, Hojsgaard D, Schmutzer T, Scholz U, Horandl E, Vogel H, Sharbel TF. 2013. Asexual genome evolution in the apomictic *Ranunculus auricomus* complex: examining the effects of hybridization and mutation accumulation. Mol Ecol 22:5908-5921.

Ruan Y, Wang H, Chen B, Wen H, Wu CI. 2020. Mutations Beget More Mutations-Rapid Evolution of Mutation Rate in Response to the Risk of Runaway Accumulation. Mol Biol Evol 37:1007-1019.

Schneider A, Charlesworth B, Eyre-Walker A, Keightley PD. 2011. A method for inferring the rate of occurrence and fitness effects of advantageous mutations. Genetics 189:1427-1437.

Schwander T. 2016. Evolution: the end of an ancient asexual scandal. Curr Biol 26:R233-235.

Snell TW, Kubanek J, Carter W, Payne AB, Kim J, Hicks MK, Stelzer C-P. 2006. A protein signal triggers sexual reproduction in *Brachionus plicatilis* (Rotifera). Marine Biology 149:763-773.

Sprouffske K, Aguilar-Rodriguez J, Sniegowski P, Wagner A. 2018. High mutation rates limit evolutionary adaptation in *Escherichia coli*. PLoS Genet 14:e1007324.

Wagner GP, Zhang J. 2011. The pleiotropic structure of the genotype-phenotype map: the evolvability of complex organisms. Nat Rev Genet 12:204-213.

Wang H-Y, Chen Y, Tong D, Ling S, Hu Z, Tao Y, Lu X, Wu C-I. 2017. Is the evolution in tumors Darwinian or non-Darwinian? National Science Review 5:15-17.

Wen H, Wang HY, He X, Wu CI. 2018. On the low reproducibility of cancer studies. National Science Review 5:619-624.

Wojciechowski MF, Hoelzer MA, Michod RE. 1989. DNA repair and the evolution of transformation in *Bacillus subtilis*. II. Role of inducible repair. Genetics 121:411-422.

Wright SI, Kalisz S, Slotte T. 2013. Evolutionary consequences of self-fertilization in plants. Proc Biol Sci 280:20130133.

Wu CI, Wang HY, Ling S, Lu X. 2016. The ecology and evolution of cancer: the ultra-microevolutionary process. Annu Rev Genet 50:347-369.

Xu X, Li G, Li C, Zhang J, Wang Q, Simmons DK, Chen X, Wijesena N, Zhu W, Wang Z, et al. 2019. Evolutionary transition between invertebrates and vertebrates via methylation reprogramming in embryogenesis. National Science Review 6:993-1003.

Zeyl C, Mizesko M, Visser JAGMd. 2001. Mutational meltdown in laboratory yeast populations. Evolution 55:909.

Zhang Y, Li Y, Li T, Shen X, Zhu T, Tao Y, Li X, Wang D, Ma Q, Hu Z, et al. 2019. Genetic load and potential mutational meltdown in cancer cell populations. Mol Biol Evol 36:541-552.

Zhou Q, Bachtrog D. 2012. Sex-specific adaptation drives early sex chromosome evolution in *Drosophila*. Science 337:341.

### Supplementary Text

#### Time to reach a steady state for Muller's ratchet

Although the approximate formulae for the fitness decline of Muller's ratchet (MR; (Haigh 1978; Gessler 1995; Gordo and Charlesworth 2000a; Gordo and Charlesworth 2000b; Etheridge, et al. 2009; Neher and Shraiman 2012)) and for the more general cases with beneficial mutations (Goyal, et al. 2012; Good, et al. 2014; Weissman and Hallatschek 2014) have been developed, the approximations are not always sufficiently accurate. Thus, we simulate a discrete-time Wright-Fisher model for the MR effect (see Methods in main text) instead. To estimate the speed of MR (i.e. the rate of mutation accumulation of asexual population), we need to confirm two things: i) the time a population need to reach a steady state while starting from a mutation-free population. (The transient effect from the initial conditions will affect the estimation of MR speed (Goyal, et al. 2012)); ii) whether the rate of mutation accumulation is constant. Here we confirmed that $10^4$ generations are generally sufficient to reach a steady state and the rate of mutation accumulation keeps steady after this equilibration time. More details are shown as follows.

As suggested by Haigh (Haigh 1978), after a particular time, mutation and selection will reach a balance in an asexual population with *N* individuals. And the frequency of individual with $i$ mutations at equilibrium will be

$$\hat{x}_i = \frac{e^{-\frac{u}{s}}(\frac{u}{s})^i}{i!}$$

where *u* is the rate of deleterious mutations and *s* is selective intensity against negative mutations (fitness loss). Furthermore, Haigh argued that it's the number ($n_{min} = N\hat{x}_0 = Ne^{-u/s}$) of least-loaded individuals to mainly determine the accumulation rate of deleterious mutations (Haigh 1978). For Muller's ratchet to work, the number of individuals in the class with the fewest mutation must be small. When $n_{min}$ becomes 0, $n_{min+1}$ becomes the new $n_{min}$ and the ratchet move a notch.

Based on computer simulations (see Methods in main text), we find the time need to reach a steady state much depends on the population size (Fig. S1 and Fig. S2). It takes ~4000 generations to reach a steady state when N is large (e.g. *N* = 10000) (Fig. S1), but less than 100 generations when *N* = 10 (Fig. S2). After the equilibration time, the number of fixed mutations, polymorphic mutations, mutations of the least-loaded individual, and the number of polymorphic mutations of the least-loaded individual will accumulate linearly. Thus, after the equilibration time, the speed of MR would be constant and then can be used to estimate the fitness decline over time. To remove the transient effects from the initial conditions, we will skip the first $10^4$ generations while simulating the dynamics of MR.

**Accuracy of approximate equations to estimate the MR's speed**

Based on Haigh's theoretical framework on linked selection, lots of researches have been done to explore MR and Hill-Robertson effect (Hill and Robertson 1966), including the approximate estimate of speed of MR (Gessler 1995; Gordo and Charlesworth 2000a; Gordo and Charlesworth 2000b; Etheridge, et al. 2009; Neher and Shraiman 2012), the shape of fitness distribution involving beneficial mutations and recombination (Goyal, et al. 2012; Good, et al. 2014; Weissman and Hallatschek 2014). Note lots of these studies is based on time-consuming simulation or approximate equations, a rigorous characterization on the interference effect is still difficult to achieve (Goyal, et al. 2012; Desai 2020). Here we test and discuss the accuracy of some approximate equations.

For example, the approximate speed of MR can be calculated by the average time it takes for the ratchet to click one notch (Gordo and Charlesworth 2000a; Gordo and Charlesworth 2000b) as follows:

$$T(N,u,s) = T_a + T_{0,x_0} + T_{x_0,1}$$

$$T_a \approx \frac{1}{s}(1 - \frac{1.6s}{u})$$

$$T_{0,x_0} = \int_0^{x_0} \frac{2N}{xG(x)} \left\{ \int_0^{x} G(y)dy \right\} dx$$

$$T_{x_0,1} = \int_{x_0}^{1} \frac{2N}{xG(x)} \left\{ \int_0^{x_0} G(y)dy \right\} dx$$

where

$$G(z) = e^{\frac{1.2Ns}{x_0} z(\frac{z}{2} - x_0)}, x_0 = e^{-\frac{u}{s}}$$

Compared to computer simulation for MR, we find the approximate equation of Gordo and Charlesworth is accurate only in some particular cases (Fig. S3 and Fig. S4). Recently, Goyal et al. (Goyal, et al. 2012) summarized that the approximate equation of Gordo and Charleworth is accurate only when $1.6 \leq \frac{u}{s} \leq 6.3$, which is consistent with our simulation results (Fig. S3 and Fig. S4). Although Goyal et al. and other researches had developed a well approximate equation for a much broader range of parameters (Etheridge, et al. 2009; Goyal, et al. 2012; Neher and Shraiman 2012; Good, et al. 2014; Weissman and Hallatschek 2014), a rigorous characterization on MR and HR is still difficult to achieve (Desai 2020).

**References**


Desai MM. 2020. Haigh (1978) and Muller's ratchet. Theor Popul Biol 133:19-20.

Etheridge AM, Pfaffelhuber P, Wakolbinger A. 2009. How often does the ratchet click? Facts, heuristics, asymptotics. LONDON MATHEMATICAL SOCIETY LECTURE NOTE SERIES:365-390.

Gessler DD. 1995. The constraints of finite size in asexual populations and the rate of the ratchet. Genet Res 66:241-253.

Good BH, Walczak AM, Neher RA, Desai MM. 2014. Genetic diversity in the interference selection limit. PLoS Genet 10:e1004222.

Gordo I, Charlesworth B. 2000a. The degeneration of asexual haploid populations and the speed of Muller's ratchet. Genetics 154:1379-1387.

Gordo I, Charlesworth B. 2000b. On the Speed of Muller’s Ratchet. Genetics 156:2137-2140.

Goyal S, Balick DJ, Jerison ER, Neher RA, Shraiman BI, Desai MM. 2012. Dynamic mutation-selection balance as an evolutionary attractor. Genetics 191:1309-1319.

Haigh J. 1978. The accumulation of deleterious genes in a population--Muller's Ratchet. Theor Popul Biol 14:251-267.

Hill WG, Robertson A. 1966. The effect of linkage on limits to artificial selection. Genet Res 8:269-294.

Neher RA, Shraiman BI. 2012. Fluctuations of fitness distributions and the rate of Muller's ratchet. Genetics 191:1283-1293.

Weissman DB, Hallatschek O. 2014. The rate of adaptation in large sexual populations with linear chromosomes. Genetics 196:1167-1183.

**Fig. S1 to Fig. S4**

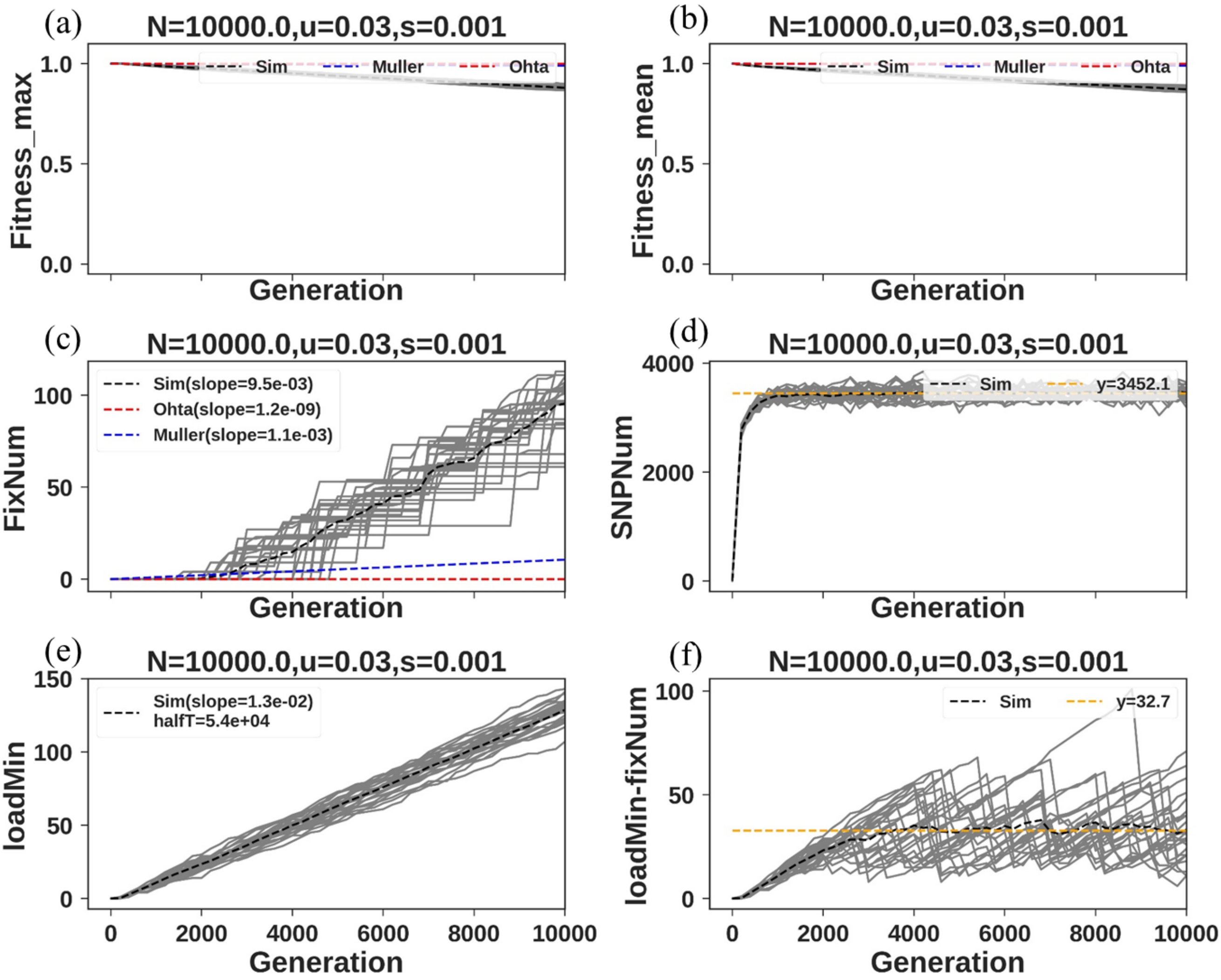


**Fig. S1 The evolutionary dynamics of MR vs. OR with deleterious mutations only (*p*=0, *q*=1)**. For all panels, $N$ = 10000, $u$ = 0.03, $s_q = s$ =0.001. The grey solid lines (25 repeats) are for MR, obtained by simulation (see Methods in main text). The black dashed line is the average of the 25 repeats. The red dashed line is the dynamics of OR over time (see main text) and the blue dashed line is the approximate equation to trace the dynamics of MR over time, obtained by Gordo and Charlesworth (Gordo and Charlesworth 2000a; Gordo and Charlesworth 2000b). (a) The fitness decline of the least-loaded individual in the population over time. (b) The decline of average fitness of the population over time. (c) The number of fixed mutations (mutations shared by all individuals in the population) over time. (d) The number of polymorphic mutations over time. (e) The number of mutations of the least-loaded individual over time. (f) The number of polymorphic mutations of the least-loaded individual (i.e. the number of all mutations of the least-load individual minus the number of fixed mutations) over time.

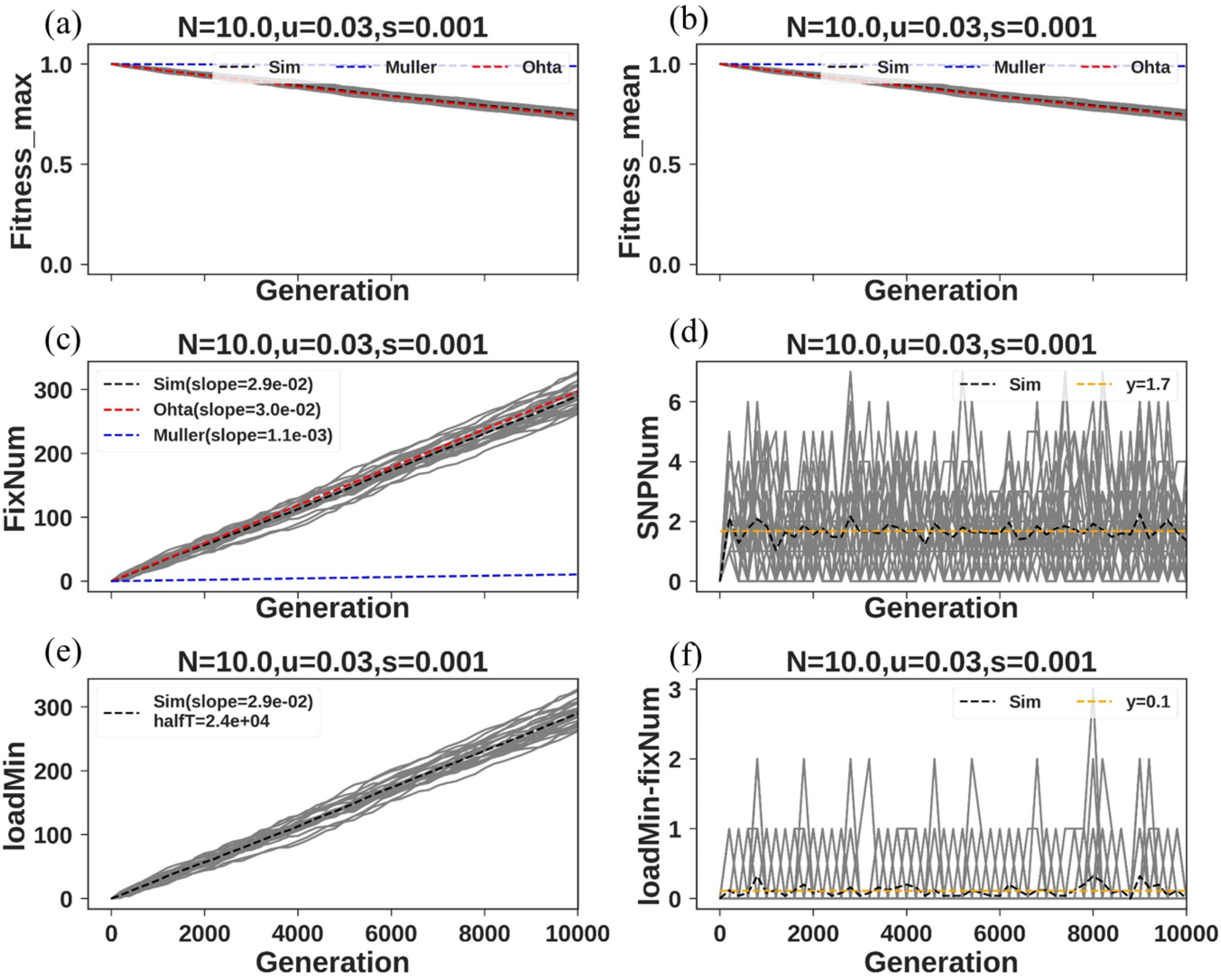


Fig. S2 same as Fig. S1 except $N = 10$.

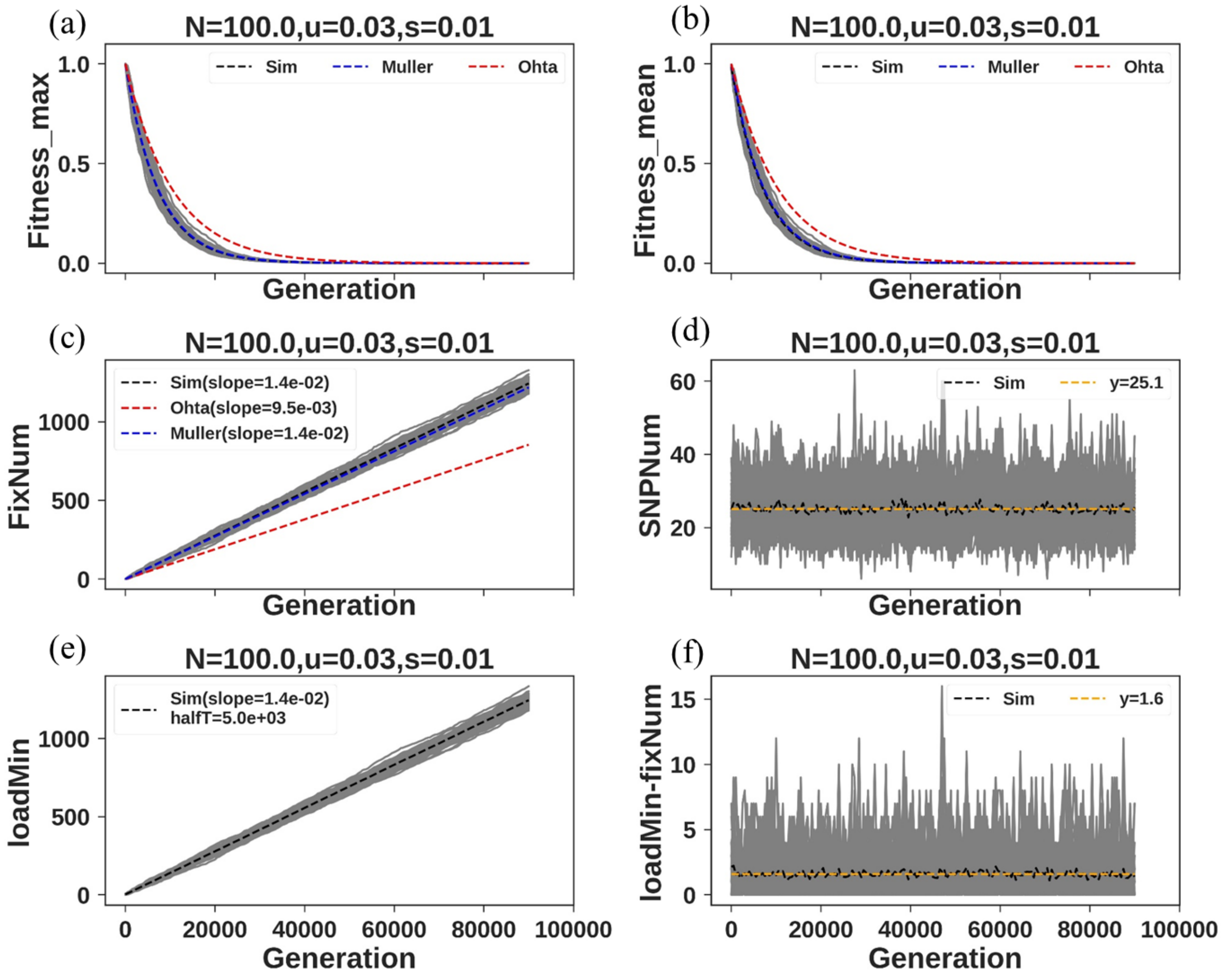


Fig. S3 same as Fig. S1 except $N$ = 100, $u$ = 0.03, $s_q$ = $s$ = 0.01.

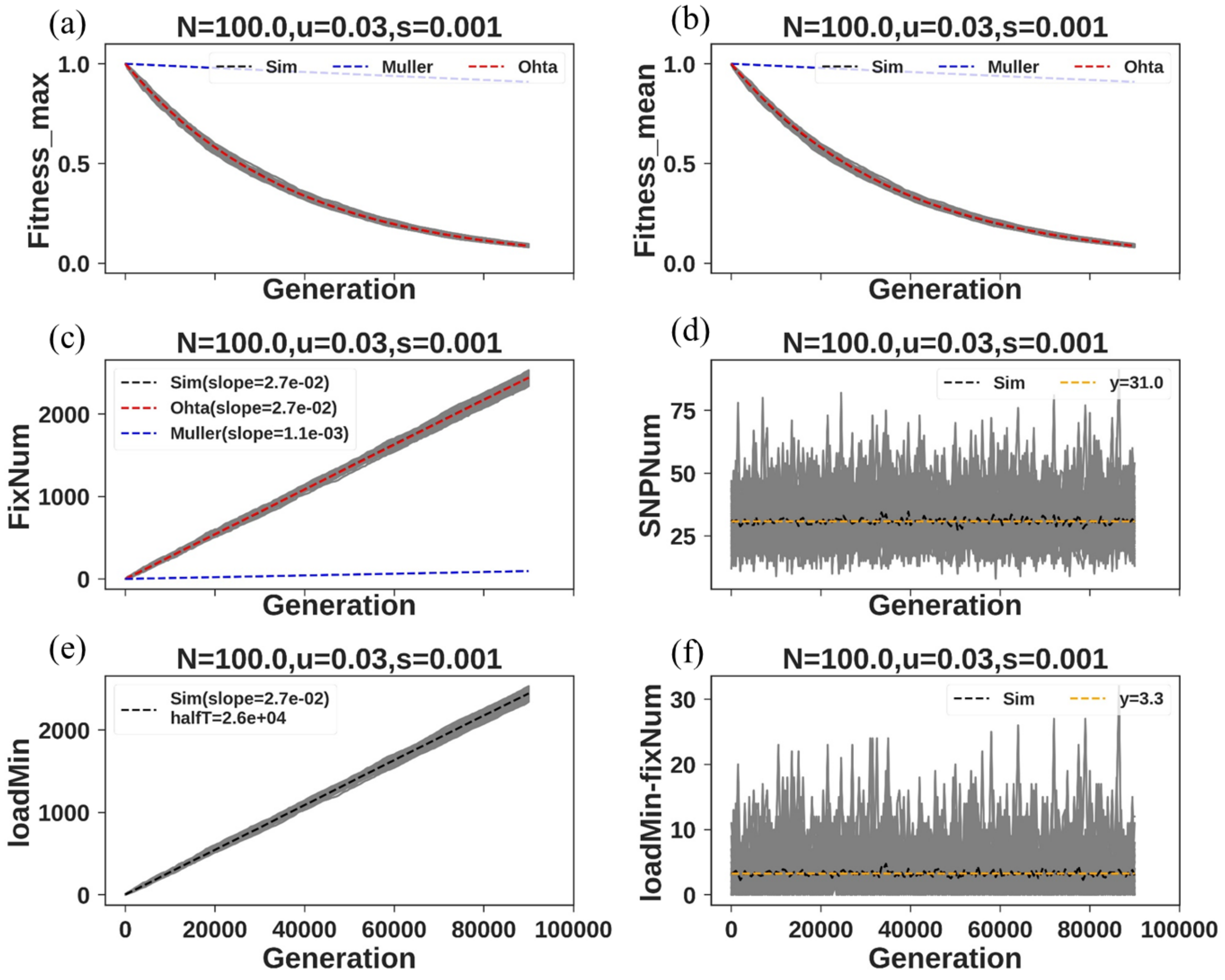


Fig. S4 same as Fig. S1 except $N$ = 100, $u$ = 0.03, $s_q$ = $s$ = 0.001.
